# The *Drosophila* fertility factor *kl-3* is linked to the Y-chromosome of the vector of Chagas’ disease *Triatoma infestans* (Hemiptera: Reduviidae) and is essential for male fertility

**DOI:** 10.1101/690370

**Authors:** Carlos H. Martins, Rodrigo S. V. P da Silva, Thalia K. Ferreira, Rafaela Paim, Grasielle C. D. Pessoa, Mauricio V. Sant’Anna, Nelder F. Gontijo, Ricardo N. Araujo, Marcos H. Pereira, A. Bernardo Carvalho, Leonardo B. Koerich

## Abstract

In many insects, the Y chromosome plays a key role in sexual determination and male fertility. The Chagas disease vector *Triatoma infestans* has 22 autosomal chromosomes and a pair of XY sex chromosomes. However, the knowledge on the Y chromosome of this species, its genetic content or its biological function, is very poor. Due to repetitive DNA, Y chromosome sequences are poorly assembled in genome projects, hindering structural and functional studies on Y-linked genes. Our group has developed many of the bioinformatic tools to identify Y-linked sequences in assembled genomes. Here, we describe the identification of a γ-dynein heavy chain linked to the Y-chromosome of *T. infestans*. This protein is orthologous to the *Drosophila melanogaster* Y-linked gene *kl-3*. In *D. melanogaster*, dyneins of the Y chromosome are known as male fertility factors and their deletion causes male infertility. We performed knockdown of the *kl-3* expression to ascertain its function in *T. infestans*. Our results showed that injection of dsKL3 reduced, significantly, the fertility of *T. infestans* males (p<0.01). The mean number of eggs laid by the control group was 35.64 eggs/couple while the kl-3 knockdown group was of 11.82 eggs/couple (five couples did not lay any eggs). Differences in eclosion rate was even more significant, with a hatching mean rate of 16.85±10.03 and 1.69±3.58 (p<0.001) for the control and the silenced groups respectively. Our results suggest that *kl-3* maintains its functional role as essential for male fertility in *T. infestans*. Hence, it seems that the Y-chromosome of *T. infestans* has a key role in male fertility. This is the first report of a *kl-3* orthologue linked to the Y chromosome of an insect species outside the diptera clade. In addition to the first report of a Y-linked gene in *T. infestans* with a role for male fertility, this finding is of great relevance for the study of the evolution of Y chromosomes and further studies that could lead to novel approaches in insect control.

## Introduction

In 1916 Calvin Bridges published his seminal paper in which he proved Morgan’s theory of Chromosomal Herdability^1^. In this same work Bridges also showed that the Y chromosome of *Drosophila melanogaster* did not determined male sex but was essential for male fertility. It was only in 1960 that Brosseau^2^ performed a series of deletion experiments showing that the *D. melanogaster* Y-chromosome contained seven fertility factors (later reduced to six fertility factors^3^), which were named as *kl-1, kl-2, kl-3, kl-5, ks-1* and *ks-2*. It took more than thirty years to find out that *kl-5* contained the coding sequence of a dynein proteins, which is responsible for the motility of flagella^4^. With the release of the *D. melanogaster* genome in 2000^5^, Carvalho and collaborators developed new strategies to identify Y-linked sequences, describing six new Y-linked genes and showing that fertility factors *kl-2* and *kl-3* harboured other dynein proteins (a x-dynein and a γ-dynein, respectively)^6,7^. Still, even in the genomic era, the study on Y-chromosome evolution and function has lagged behind. The Y chromosome is heterochromatic in most species, which makes it difficult to identify Y-linked sequences in many genome studies^8,9^. Even now, with accessible sequencing technologies, the most studied Y chromosomes are those of mammals (specially humans, chimpanzees and mice) and *Drosophila*^6,7,10–17^. Recent studies proposed new approaches to use the power of new genome sequencing technologies (NGS) to boost the identification of Y-linked sequences in new genomes. In the first study, Hall and collaborators^18^ performed Illumina DNA sequencing of males and females of *Anopheles* mosquitoes, and were able to identify six new Y-linked genes in these insects. This method was also used to describe the male sex determination gene in *Aedes aegypti*^19^. In the second study, Carvalho and Clark^20^ used sequences from female DNA to find specific male sequences in the assembled *Drosophila virilis* and human genomes. They were able to identify four new Y-linked genes in *D. virilis* and 300 kb of previously unidentified sequences on the human Y chromosome. Insects have a variety of sex chromosome systems, ranging from total absence of sex chromosomes to X0, X^n^Y, ZW and traditional XY. Therefore, the study of genomes of non-model insects, such as insects’ vectors of disease, could provide valuable data to understand sex-chromosome evolution and promote the expansion on the knowledge of insect biology.

Chagas disease is one of the most important parasitic infection in Latin America and more than 12 million people are infected by *Trypanosoma cruzi* (the protozoan agent of Chagas’ disease)^21^. *Triatoma infestans* is the most important vector species in the southern cone area of South America and with the effort of the Southern Cone Initiative (which had the main goal of interrupting the *T. cruzi* transmission using chemical insecticides to eliminate *T. infestans* populations)^22^ its populations were highly reduced. However, *T. infestans* persists as domestic and sylvatic populations in several areas of the Gran Chaco region from Argentina, Bolivia and Paraguay and parts of the highland valleys of Bolivia^23^ and studies suggests that high genetic polymorphism could be correlated to *T. infestans* resistance. Despite their medical importance, the research on triatomine genetics is almost non-existent and only the genome of *Rhodnius prolixus* is available so far. Cytogenetical studies suggests that Andean and non-Andean populations of *T. infestans* have a significant difference in genome size (1.8Gbp and 1.1Gbp respectively), despite a constant number of diploid chromosomes (10A + XY), which suggests a high variability in heterochromatin^24,25^.

In 2016 we described nine new Y-linked genes in *R. prolixus*^26^. At that time, we could only speculate on triatomine Y-chromosome role and on the origin of these chromosomes. More than that, we pointed out the need of new triatomine genomes for further evolutionary studies. Here we describe a thorough analysis of the *T. infestans* Y-chromosome. Differently from the *R. prolixus* Y-chromosome (in which we were not able to find single copy genes), the *T. infestans* Y-chromosome harbors a single copy γ-dynein protein and we provide evidence that, as in *D. melanogaster*, this protein is essential for *T. infestans* male fertility.

## Results and Discussion

Chagas disease is one of the most important parasitic infection in Latin America and more than 12 million people are infected by *Trypanosoma cruzi* (the protozoan agent of Chagas’ disease)^21^. Despite its relevance as vector of Chagas’ disease, triatomine genetics has been neglected for many years. The recent sequencing of *R. prolixus* and the effort to sequence *T. infestans* (unpublished) has facilitated some genetic studies, mainly in functional genomics using RNAi^27,28^. Still, a lot is missing in our knowledge of gene function, chromosome organization and evolution. Y-chromosomes are involved in major biological phenomena such as sex determination and male fertility. They remain largely uncharacterized because of their high repeat content which precludes sequence assembly into large and easily studied contigs. Using the approach proposed by Carvalho and Clark (called YGS) we identified many Y-linked sequences and found a single copy γ-dynein that is orthologous to the *D. melanogaster kl-3*.

### Identification of Y-linked genes in *Triatoma infestans*

*Triatoma infestans* genome was assembled into 4,865 scaffolds that cover 1.1Gb (∼90% of the genome). The number scaffolds that cover 50% of the genome (N50) is 224, the N50 scaffold size is 1.1Mb. The YGS program^20^ determines the percentage of *k-mers* per scaffold that are unique to the male genome. Figure 1 (panel A) shows the distribution of scaffolds by size (number of valid *k-mers*) per percentage of unique male sequences. In our initial analysis, the autosomal or X sequences are shifted to the right of the graph (close to the 5% point of single male sequences), while there is no 100% male sequence. Carvalho *et al*.^20^ pointed out that such results suggest, respectively, low coverage of female DNA sequencing and slight contamination of male DNA in female DNA samples. As suggested by Carvalho and cols.^20^ we considered all scaffolds with more than 30% as putative Y-linked sequences. This cut-off returned 334 scaffolds (∼5.8Mbp of sequences) as candidates for Y-linkage. Nearly 30% of the sequences (1.98Mbp) were constituted of gaps, and initial filtering eliminated 217kbp of satellite DNA and 8.4kbp of ribosomal DNA (rDNA). From the remaining sequences, initial blast searches identified 3,146 possible coding sequences, from which 2,637 (83.8%) were identified as transposable elements. We selected 12 scaffolds containing putative coding sequences to test linkage to Y as proof that the proposed method works (Figure 1, panel B).

**Figure 1.**
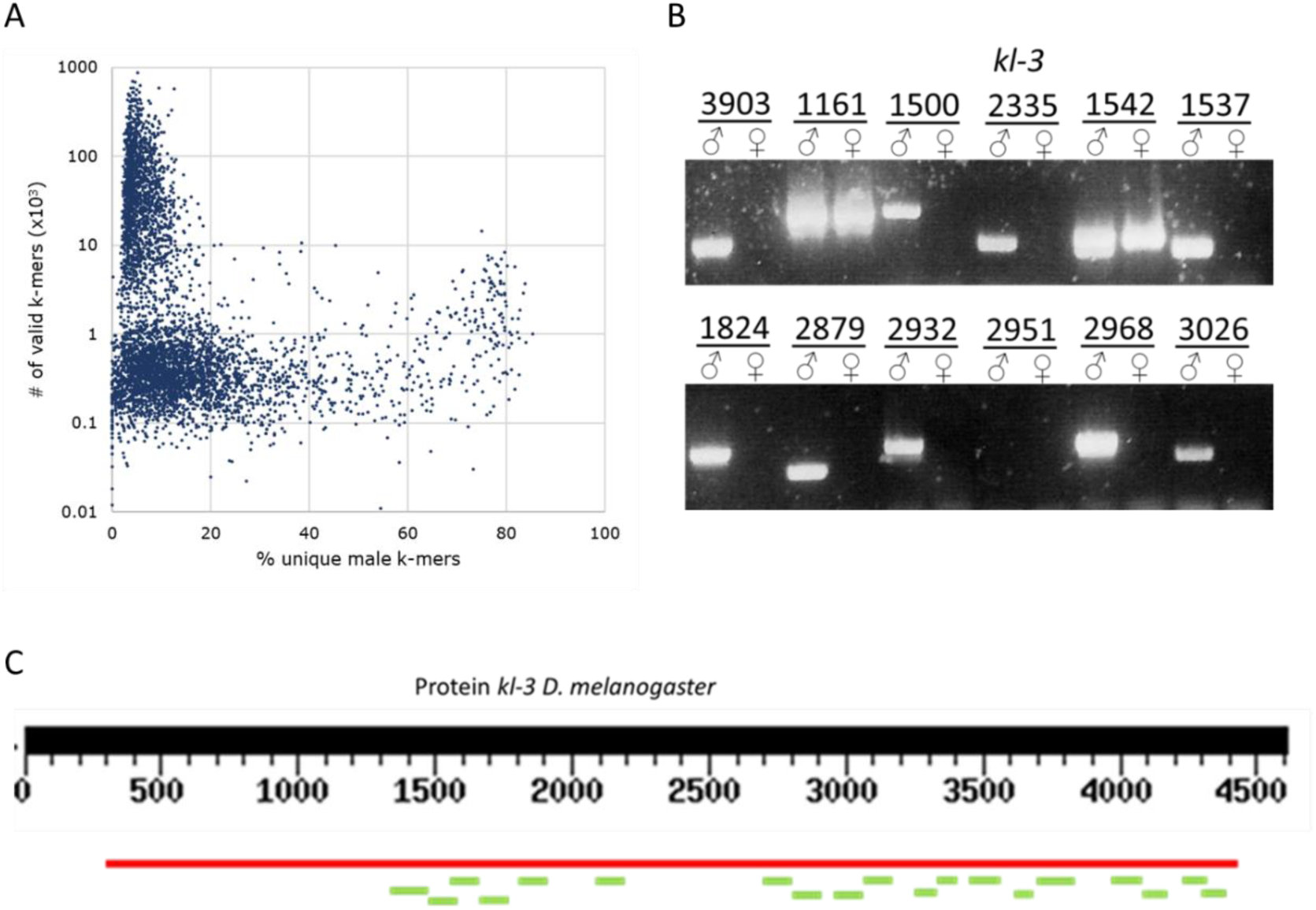
Identification of Y-linked sequences of *Triatoma infestans* and *kl-3* annotation. **A)** *T. infestans* scaffolds (blue dots) were evaluated according to the number of single-stranded sequences (*k-mers*) that lack homology to female sequences (% unique male sequences). For a preliminary analysis, we consider scaffolds with more than 75% unique sequences as Y-linked candidates. **B)** Linkage assay was performed by PCR for male and female DNA. The presence of amplification in male with absence of amplification for female samples indicates linkage to the Y chromosome. Sequence 2335 contains an exon of the *kl-3* gene. **C)** tBLASTn search using the *D. melanogaster kl-3* gene as a template suggests the existence of two γ-dynein genes in *T. infestans*. One of the genes is found in scaffold 603 (red bar) which possesses only 4% of unique male sequences, suggesting linkage to an autosomal chromosome or to the X chromosome. The second copy is dispersed in several candidate Y chromosome scaffolds (green bars).

The remaining 509 putative coding sequences were grouped into 390 clusters. Virtually all clusters found were composed of multicopy genes (those clusters contained the following *R. prolixus* Y-linked genes: *met-Y, znf-Y1, znf-Y2*; *rpr-Y2* and *rpr-Y3*), while only three clusters were composed of single copy genes. From those, two showed similarities with RNA-polymerases while a third cluster (composed of 19 sequences) showed high similarities with γ-dynein heavy chain proteins. It is important to make a few considerations here. Many studies using genomics to identify Y-linked genes have shown the difficulties on studying heterochromatic regions^6,7,18–20,29^. Genome assemblers based on short read sequences are not capable to assemble repetitive regions, such as satellite rDNA. Hence, in most genome projects, Y-linked scaffolds are usually composed of short contigs or are packed with gaps (in our case, gaps enclose nearly 30% of the scaffold sequences)^6^. Also, heterochromatic sequences are well known for its richness in transposable elements (TEs), which creates an environment that facilitates the increase in copies of heterochromatic genes^8,30,31^. Here we found that from 3,146 putative coding sequences, 2,637 are of TEs and at least 487 are multicopy genes. A total of 22 coding sequences proved to be unique (single copy) and 19 of them showed similarities with a single γ-dynein protein. Carvalho and cols. were the first to point out that, due to the nature of Y genome assembly, Y-linked genes are usually scattered, and incomplete, in assembled genomes, creating a signature of Y-linked genes^6^. Figure 1 (panel C) shows the alignment of these 19 sequences (green bars) with the *D. melanogaster* γ-dynein *kl-3* (Figure 1, panel C) in which the scattered pattern expected for Y-linked genes is clear. The same figure also shows a great number of gaps in the Y-linked gene, and an autosomal paralog in scaffold 603 (red bar) that has only 4% of unique male *k-mers*. We are working in filling all the gaps by standard Sanger sequencing; however, this process is laborious and remains unfinished.

### The Y-linked *γ*-Dynein of *Triatoma infestans* is orthologous to the fertility factor *kl-3* of *Drosophila melanogaster*

Although BLAST results suggested that the Y-linked γ-dynein was similar the *D. melanogaster* fertility factor *kl-3*, it was only with an evolutionary study that we could ascertain this relationship. Indeed, using the *D. melanogaster kl-3* paralogous gene to ascertain ortholog status, phylogeny strongly suggests that the Y-linked γ-dynein is orthologous to *kl-3* (Figure 2). *Kl-3* is not exclusive of *D. melanogaster* and *T. infestans* and can be found in many insect species. However, in most species with sequenced genomes, *kl-3* is an autosomal gene. In fact, we do not have empirical evidence for *kl-3* linkage in every species. However, the fact that in all these species (besides *Drosophila* and *T. infestans*) *kl-3*, a ∼15kbp gene, is easily found complete in large scaffolds, is a strong evidence for autosomal linkage. Interestingly, we could not find (even in the read archives) the *kl-3* in *R. prolixus* (although we have found its paralogous gene CG9492). This brings the question on when *kl-3* was lost in *R. prolixus and* answering this question may bring interesting data on triatomine Y chromosome evolution. However, to answer such question, new genomes on the Rhodinii and Triatominii tribes are necessary.

**Figure 2.**
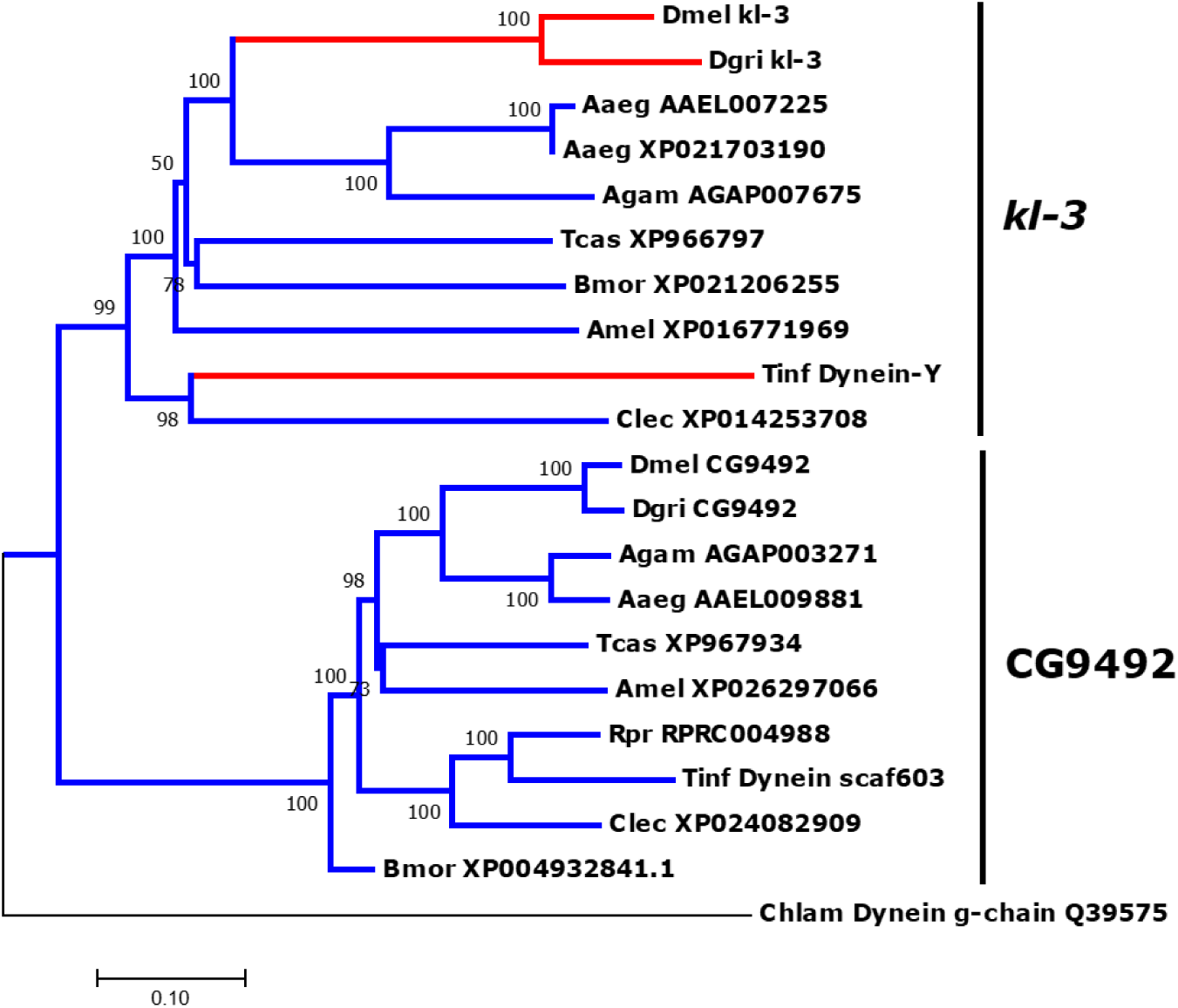
Evolutionary analysis of *Triatoma infestans kl-3*. The evolutionary history of the *kl-3* and its paralogous gene CG9492 was inferred using the Neighbor-Joining method. Red lines show lineages in wich *kl-3* is linked to the Y-chromosome. Blue lines show lineages in which the referred gene is putatively autosomal. Sequence accession numbers are shown in each branch after species abbreviation. Species abbreviations are: Dmel = *Drosophila melanogaster*; Dgri = *D. grimshawii*; Aaeg = *Aedes aegypti*; Agam=*Anopheles gambiae*; Tcas = *Tribolium castaneum*; Bmor = *Bombyx mori;* Amel = *Apis mellifera*; Rpr = *Rhodnius prolixus*; Tinf=*Triatoma infestans*; Clec=*Cimex lectularius.* Evolutionary analyses were conducted in MEGA.

### Silencing the expression of *Triatoma infestans kl-3* causes male sterility

After the ascertain of the Y-linked γ-dynein status as orthologous to *kl-3*, we questioned ourselves if this gene was essential for male fertility. We first quantified the *kl-3* mRNA expression during the insect development (Figure 3, panel A). We observed that *kl-3* mRNA starts to be expressed in very low levels in 4^th^ instars. The expression levels increase significantly in 5^th^ instar males (when testis development is observable) and maintain similar levels in adult males, with specific expression in testis. Interference RNA has been successfully applied to triatomine functional genomics for more than ten years^28,32^, and recent studies suggests that gene knockdown are epigenetically heritable in *R. prolixus*^27^. To understand if *T. infestans kl-3* is essential for male fertility, we injected 5^th^ instar males with 300ng of double stranded RNA targeting the *kl-3* mRNA (dsKL3). After moulting, dsKL3 treated animals were used to form couples with virgin females, and egg laying was accompanied for five weeks. Our results show that *kl-3* knockdown reduced the mean oviposition from 35.64±21.22 to less than 11.82±17.55 eggs laid per couple (Figure 1, panel B). While the control group presents a normal distribution of oviposition, the treated group presented a clear abnormal distribution (with many females not laying any eggs). Kolmogorov-Smirnov tests showed that the 3-fold reduction in oviposition is statistically significant (p<0.01). When the number of individuals in the progeny was evaluated, the differences were even more significant (p<0.001), with a reduction in the number of offspring (Figure 1, panel C) from 16.85±10.03 (control group) to 1.69±3.58 (dsKL3 group).

**Figure 3.**
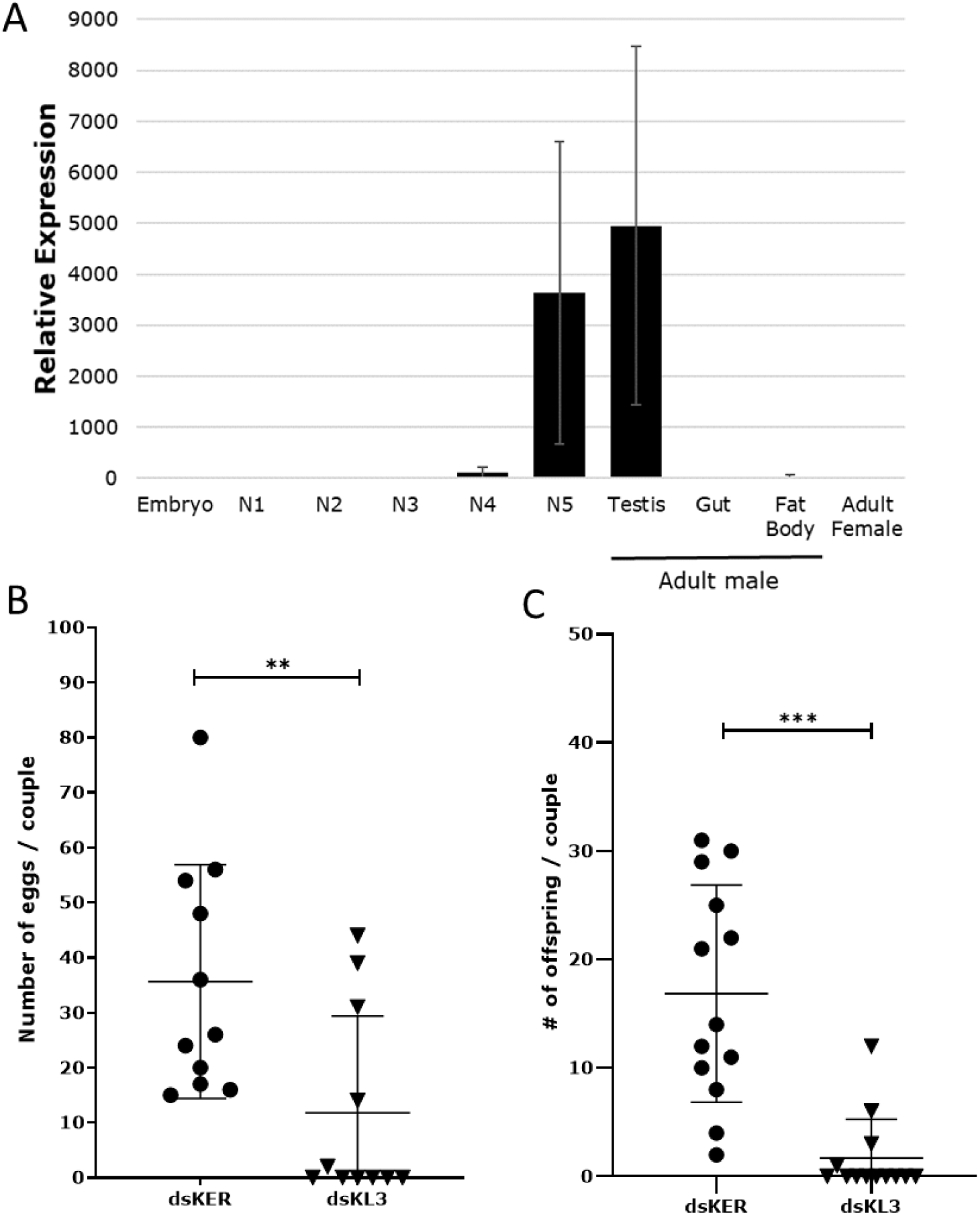
Functional analysis of *Triatoma infestans kl-3.* **A)** Expression pattern of *kl-3* in different development stages and adult tissues of *T. infestans.* **B)** Number of eggs laid per couple of *kl-3* knockdown (dsKL3) and control (dsKER) males. Fifth instar males were injected with 1µg of dsRNA and couples were formed with virgin females **C)** Number of offspring per couple of *kl-3* knockdown (dsKL3) and control (dsKER) males.

In *D. melanogaster*, knockout of *kl-3* causes sperm immobility. Immediately, a question whether females were in fact inseminated and spermatozoa was immotile, was raised. Indeed, spermatheca visualization in optical microscope showed that all females, from both groups, were fertilized. However, there is no mention on the literature about sperm motion in triatomines, and there is no protocol for such observations. Hence, the evidences shown from now on are based on exploratory approaches and with no quantitative procedures. In our slide preparations, we could not find sperm motility in many samples from the control group. In those cases, were motility was observed, it took up to ten minutes to observe some activity. For the *kl-3* knockdown group, the number of slides were motility was observed was even lower, and it took up to 20 minutes to observe any activity. However, we could not observe any correlation between egg laying and sperm motility in dsKL3 treated females. As stated before, our methodology was purely based in intuition and we are now preparing better protocols and searching for objective quantitative methods to better understand how *kl-3* knockdown impairs sperm motility. Nonetheless, this is the first report of sperm motility in triatomines. One important issue that remains to be answered is the reason why dsKL3 treated females did not laid eggs, since in *R. prolixus* virgin females lay eggs normally. There is nor report if *T. infestans* females lay eggs normally and we are performing experiments to answer that question.

## Conclusions

The elusive nature of Y-chromosomes hindered many studies on its genetics and evolution for decades. With the accessibility to genome sequencing, we are now living a gold era on Y-chromosome genomics, even in non-model species. Triatomine insects are vectors of one most important parasitic disease in Latin America, the Chagas’ disease, and *Triatoma infestans* is the most triatomine relevant vector in the souther cone region of South America. Despite its relevance in public health, triatomine genetics has lagged behind. As a result, until now, we had no idea on the role of triatomine Y-chromosomes on sex determination and male fertility.

In our study we found the first Y-linked gene on the *Triatoma infestans* Y-chromosome. This Y-linked gene is a γ-dynein protein, orthologous to the Y-chromosome fertility factor *kl-3* of *Drosophila melanogaster*. This is the first time that *kl-3* is found linked to the Y-chromosome outside the Diptera clade. Through the use of gene silencing by interference RNA, we also provide that *kl-3* is essential for *T. infestans* male fertility. Thus, this is the first report providing evidence that *T. infestans* Y-chromosome has a role on male fertility. Our findings do not only provide new knowledge on triatomine biology, but also could offer some insights on triatomine Y-chromosome evolution. Moreover, we make available some valuable data on the scarce knowledge on triatomine male biology. Further triatomine genomes could provide even more information on the origin and evolution of triatomine Y-chromosome, ascertain if this chromosome is essential for sex-determination and provide new tools for biology and vector control studies.

## Acknowledgments

This research is part of the Brazilian research program Institutos Nacionais de Ciência e Tecnologia – Entomologia Molecular.

## Author contributions

We declare that this work was done by the authors named in this article and all liabilities pertaining to claims relating to the content of this article will be borne by the authors. All authors wrote, reviewed and approved the manuscript including figures and tables. L.B.K. conceived the study. R.S.V.P.S., L.B.K and A.B.C identified Y-linked sequences and performed the gene annotation process. R.S.V.P.S. performed Y-linkage confirmation tests, C.H.M. and R.S.V.P.S. performed the expression analysis. C.H.M., T.K.F and R.P. performed functional experiments. C.H.M., R.N.A., M.R.V.S., N.F.G., M.H.P., G.D.P. and L.B.K evaluated and discussed the results. L.B.K. edited the manuscript.

## Additional information

### Data availability

All sequences and annotation tables will be freely available in the GenBank as soon as the manuscript describing the *T. infestans* genome is published.

## Material and Methods

### Insects and genomic data

Genomic lineage of *Triatoma infestans* was obtained from a colony maintained at FIOCRUZ-RJ from which insects were originally collected from the municipality of Mato Queimado, Rio Grande do Sul (Brazil). The manuscript describing the *T. infestans* genome sequencing strategy and annotation is under preparation. Briefly, genomic DNA was purified from adult males and from 5^th^ instar females and used to build male and female libraries. Each library was Illumina sequenced for a total depth of 50x and assembled with AllPathsLG^33^ into ∼4,000 scaffolds, covering 1.1Gbp (95%) of the genome. For molecular biology experiments, we used *T. infestans* maintained at our laboratory (Federal University of Minas Gerais) that was initiated with insects from populations of different regions of Brazil. All insects were maintained in a controlled room temperature (34°C) and humidity (60%), and feed in chickens every 14 days.

### Identification and annotation of Y-linked genes

The identification of Y-linked candidates was performed following the YGS method^20^, using male reads to validate the results. All *T. infestans* scaffolds identified as Y chromosome candidates were softmasked for simple repetitive sequences using RepeatMasker software and the blast masking tool against a repetitive elements database (which included rDNA, transposable elements and multicopy genes, such as histones). Masked scaffolds were then blasted against different databases (Non-redundant Protein, Ref-Seq, SwissProt and *R. prolixus* proteins) in order to search for gene coding regions. A third Blast search against bacterial proteins was used to further filter our results (nucleotide or aminoacid identities above 95%). Alignments produced from at least one of the databases were considered as putative Y-linked genes. Putative genes were ordered according to the evidence, which were: 1) alignment with a conserved known gene from Ref-Seq or NR databases; 2) alignment with a conserved hypothetical gene from Ref-Seq database; 3) alignment with a *R. prolixus* annotated gene; and 4) alignment with a unconserved hypothetical gene from Ref-Seq or NR databases. Putative Y-linked genes were then blasted against the *T. infestans* genome to identify single-copy genes, recent duplications and multicopy genes. Y-linkage was confirmed by PCR as described elsewhere^6,7,26,29,34^. Confirmed Y-linked genes were then annotated with GeneWise to identify truncated genes and to define strategies for complete gene annotation. All Y-linked genes without described function were blasted against Protein Family of Domains database (PFam) and Eukariote Conserved Orthologous Database

### Molecular biology methods

Males and virgin female *T. infestans* genomic DNA were isolated using DNeasy Blood & Tissue Kit (Qiagen, cat# 69504). PCR for the detection of Y-linkage was performed with GoTaq® Hot Start Polymerase (Promega, cat# M5005) and primers were designed to target exons (for gene Y-linkage tests) or elsewhere in the scaffold (for scaffold Y-linkage tests). RNA was isolated with TRIzol® Reagent (Invitrogen, cat# 15596–018), following manufacturer instruction from pools of five individuals and for different developmental stages and tissues (embryos, all five instar stages, female whole body, male testis, male gut, male fat body and male carcass). cDNA was synthesized using High-Capacity cDNA Reverse Transcription Kit (Life Technologies, cat# 4368814). *Kl-3* expression was evaluated by qRT-PCR using the specific primers (Ti_kl3_qF1 5’-ACCTACCCCAGCTAATTTTCAC-3’ and Ti_kl3_qR1 5’-GACATTCCTCGCCTTTAATTGAC-3’) using the *T. infestans* actin gene 18S as control (Ti_18S_F 5’-TTGGGGCTTGCAATTGTTCC-3’ and Ti_18S_R 5’-TACAAAGGGCAGGGACGTAATC-3’). Rapid Amplification of cDNA ends (Invitrogen, cat# 18373–019 and 18374–058) and RT-PCR (Invitrogen, cat# 12574– 035) were performed for gene annotation and nucleotide sequencing correction. All PCR products were Sanger sequenced at Macrogen (Korea).

### Functional analysis through knockdown experiments

Functional analysis was carried out using gene silencing (knockdown) by interference RNA (RNAi) using standard methods patronized by our group^27,32^. Basically, double stranded RNA for *kl-3* (dsKL3) and the control gene keratine (dsKER)^27^ were synthesized using the MegaScript High Yeld Transcription Kit (Ambion - USA) according to the manufacturer’s instructions (primers Ti_kl3_t7F1 5’-TAATACGACTCACTATAGGGGTTTTGTCCTGGAATTATTG-3’ and Ti_kl3_t7R1 5’-TAATACGACTCACTATAGGGACATCACCTGTAAGAAATAC-3’; product size 533bp). After suspension of dsRNA (1µg/ul) in sterile saline solution, 300ng of each dsRNA was inoculated in 5^th^ instar males for the respective treatment groups. Each experimental group (dsKL3 and dsKER) was composed of ten insects that were fed weekly in mice (hairless) after dsRNA injection. Molting and death were also observed weekly. Emerging adults were then transferred to flasks containing a virgin adult female. Couples were feed weekly on mice (hairless) and egg laying was annotated for five weeks after couple formation. Egg hatching was accounted for two extra weeks. Males were dissected on the fifth week post couple formation to measure RNA expression levels (qRT-PCR), while females were dissected to confirm insemination. Female spermatheca was dissected in sterile saline solution (pH 7.0) and transferred to a microscope slide with 5µl of sterile PBS (pH 6.8). Spermatheca were then sheared with the use of sterile needles and then a micro slide laid upon the mixture. A final pressure in the microslide with the tip of a pen was applied to spread the material. Each slide was visualized in optical microscopes (400x magnification) for up to 30 minutes or until resection.

### Evolutionary tests and statistical analysis

Dynein amino acid sequences sequences from other organisms were obtained at NCBI and aligned with Muscle^35^. Evolutionary analyses were conducted in MEGA7^36^. The evolutionary history was inferred using the Neighbor-Joining method (10.000 replicates; pairwise deletion)^37^. The evolutionary distances were computed using the Poisson correction method and are in the units of the number of amino acid substitutions per site. All accession numbers are shown in the respective figures. Statistical analysis was performed in GraphPad Prism 8.1.2 software (GraphPad Software Inc.)

